# The metabolome of individuals with knee osteoarthritis is influenced by 18-months of an exercise and weight loss intervention and sex: the IDEA trial

**DOI:** 10.1101/2024.02.15.580523

**Authors:** Hope D. Welhaven, Avery H. Wefley, Brian Bothner, Stephen P. Messier, Richard F. Loeser, Ronald K. June

## Abstract

**Objective:** The Intensive Diet and Exercise for Arthritis (IDEA) trial was conducted to evaluate the effects of diet and exercise on osteoarthritis (OA), the most prevalent form of arthritis. Various risk factors, such as obesity and sex, contribute to the debilitating nature of OA. While diet and exercise are known to improve OA symptoms, cellular and molecular mechanisms underlying these interventions, as well as effects of participant sex, remain elusive.

**Methods:** Serum was obtained at three timepoints from IDEA participants assigned to groups of diet, exercise, or combined diet and exercise (n=10 per group). All serum metabolites were extracted and analyzed via liquid chromatography-mass spectrometry combined with metabolomic profiling. Extracted serum was pooled and fragmentation patterns were analyzed to identify metabolites that statistically differentially regulated between groups.

**Results:** Changes in metabolism across male and female IDEA participants after 18-months of diet, exercise, and combined diet and excise intervention mapped to lipid, amino acid, carbohydrate, vitamin, and matrix metabolism. The diverse metabolic landscape detected across IDEA participants shows that intervention type impacts the serum metabolome of individuals with OA in distinct patterns. Moreover, differences in the serum metabolome corresponded with participant sex.

**Conclusions:** These findings suggest that intensive weight loss among male and female subjects offers potential metabolic benefits for individuals with knee OA. This provides a deeper understanding of dysregulation occurring during OA development that may pave the way for improved interventions, treatments, and quality of life of those impacted by this disease.

## Introduction

Obesity is the most modifiable risk factor and is a known accelerant of osteoarthritis (OA)(1, 2). Weight loss via diet and exercise improves symptomatic knee OA and is recommended for overweight and obese patients with OA(3, 4). However, the cellular and molecular mechanisms underlying weight-loss-related symptom improvement remain elusive. OA affects both males and females, but knee OA is more severe and prevalent among females(5, 6). Known female-specific OA risk factors include genetics, anatomy, and increased likelihood to sustain a traumatic knee injury compared to male counterparts(7, 8), although sexual dimorphism at the molecular and cellular levels remains unclear.

Metabolomics, the study of small molecule intermediates called metabolites, can provide insight into OA pathogenesis, patient risk factors, and metabolic perturbations caused by diet and exercise interventions. This powerful tool has been applied in other musculoskeletal studies to analyze the metabolism of various fluids and tissues (e.g., synovial fluid) during disease progression(9–13) and after mechanical loading(14–16). NMR metabolomics provided insight into urine signatures in the multi-faceted Intensive Diet and Exercise for Arthritis (IDEA) trial (ref 17 belongs here as well as below). The IDEA trial was conducted to assess the effects of weight loss, via diet and exercise, on primary and secondary outcomes in OA participants. Using NMR (Nuclear Magnetic Resonance) metabolomics, dysregulated amino acid and lipid metabolism were found in a subset of IDEA urine samples from participants with radiographic progression compared to non-progressors (17). While this study provided insight into disease progression based on urine profiles, further investigation is required to better understand current methods of weight loss, the role of obesity and sex, and underlying molecular mechanisms driving OA.

Therefore, the objectives of this study were to (1) compare serum global metabolomic profiles to analyze changes in metabolism over the course of 18-months of an exercise and weight loss intervention, (2) illuminate metabolic pathways influenced by different types of interventions, and (3) identify signatures that are sex dependent.

## Materials and Methods

### Study Design, Interventions, and Participants

The participants in this study were a subset of those enrolled in the IDEA trial. In total, 30 participants were randomly selected (n=30) with 10 participants per intervention group (Diet, D: n = 10, Exercise, E: n = 10, Diet and Exercise, DE: n = 10). Participants included in this study were matched for age, sex, and BMI (Table 1). Race and ethnicity were not provided for this study, thus were not assessed.

**Table 1.** Participant information for each intervention type. Baseline BMI and age are detailed by group and broken down by sex. Data are presented as mean ± standard deviation from the mean.

|  | Exercise (E) | Diet (D) | Diet & Exercise (D+E) |
| --- | --- | --- | --- |
| BMI | 33.14 ± 2.03 | 33.39 ± 2.20 | 32.93 ± 2.83 |
| BMI Female | 32.75 ± 2.52 | 32.75 ± 2.52 | 33.75 ± 3.27 |
| BMI Male | 33.53 ± 1.58 | 34.02 ± 1.89 | 32.12 ± 2.37 |
| Age | 65.99 ± 4.21 | 65.78 ± 4.34 | 65.89 ± 1.79 |
| Age Female | 65.80 ± 5.26 | 65.80 ± 5.26 | 65.65 ± 0.62 |
| Age Male | 66.19 ± 3.50 | 65.76 ± 3.86 | 66.11 ± 2.59 |

The IDEA trial was conducted at Wake Forest University and Wake Forest School of Medicine between July 2006 and November 2011. This trial was a single-blind, 18-month, randomized controlled trial to determine whether a ≥10% loss in body weight induced by different intervention types would improve primary (e.g., knee joint compression forces and IL-6 levels) and secondary clinical outcomes (e.g., pain, function, mobility). Participants (n=454) were assigned to one of three interventions: diet (D), exercise (E), or diet and exercise (D+E). Additional details about intervention types, radiographic measurements, participant inclusion criteria, and study design are provided in the initial study reports(3, 4).

Diet parameters for D and D+E participants included up to 2 meal-replacement shakes per day, and a third meal selected from a weekly menu plan composed of traditional foods within 500-760 kcals each. Participants were allowed to have snacks (snack bar, fruit, or vegetables) that were 100-120 kcal each. All meal plans were developed and overseen by staff to assure macronutrient-balanced energy intake. Exercise parameters included three sixty-minute sessions per week for 18 months. For the first six months, the sixty-minute sessions were center-based and consisted of an aerobic phase (i.e., walking) for 15-minutes, a strength training phase for 20-minutes, another aerobic phase, and a cool-down phase for 10-minutes. For the remainder of the study, participants had the option of continuing to exercise at the center, transitioning to a home-based exercise routine, or using a combination of center and home-based exercise(3).

### Serum Sampling, Extraction, and Metabolite Profiling

Blood samples were collected from participants following a 10-hour fast at three different time points: baseline, 6-months, and 18-months(4). At each time point, samples were collected, and serum was stored at -80°C until extraction. To extract serum metabolites, 100 uL aliquots were centrifuged at 500xg for 5 minutes to remove cells and debris. Next, supernatant was transferred to a fresh mass-spectrometry grade microcentrifuge tube, 500 uL of cold acetone were added, samples were vortexed vigorously, and chilled overnight at -80°C to precipitate proteins. The following day, serum samples were vortexed again, centrifuged at 16,100xg for 10 minutes, and supernatant was evaporated by vacuum concentration. Once dry, metabolites were resuspended with 1:1 acetonitrile:water.

The extracted serum samples (n=90) were analyzed using liquid chromatography-mass spectrometry (LC-MS). Samples were analyzed in positive mode using an Agilent 1290 LC coupled through an electrospray ionization source to an Agilent Quadrupole Time of Flight (Q-TOF) mass spectrometer. Ions were separated using a Cogent Diamond Hydride HILIC chromatography column (2.2 µM, 120 Å, 150 mm x 2.1 mm, MicroSolv Leland, NC, United States) at a flow rate of 0.400 uL/min. For quality control purposes, the order of sample injection was randomized, 5 uL of each sample was injected, and blank samples containing neat 1:1 acetonitrile:water were injected every 10 samples.

LC-MS data consisted of mass-to-charge ratios (m/z), metabolite abundances, and retention times which were processed using Agilent Masshunter Qualitative Analysis software and MSConvert(18). Data were then exported and converted using XCMS(19). Using in-house standardized procedures(20), MetaboAnalyst (version 5.0) was used to perform a suite of statistical analyses to visualize data and distinguish subsets of metabolite features. These included: hierarchical clustering analysis (HCA), principal component analysis (PCA), Partial Least Squares-Discriminant Analysis (PLS-DA), ANOVA, volcano plot analysis, fold change, and student’s t-tests. Additionally, fold change, volcano plot, and median intensity heatmap analyses were used to find metabolite features that are co-regulated and dysregulated between groups. Populations of metabolites distinguished by these three analyses underwent pathway enrichment analyses using the Mummichog algorithm within MetaboAnalyst to map clusters of key metabolites to cellular pathways. Significance was determined using FDR-corrected p-values with an *a priori* threshold of p < 0.05.

### Metabolite Identification through Pooled Analysis

Additionally, 10 pooled samples were created by combining original extracts. For each pool, 10 uL from 5 randomly selected participant samples were combined. This process was repeated for all pools (n = 10). Pooled samples were then subjected to liquid chromatography tandem mass spectrometry (LC-MS/MS) for metabolite identification purposes. Pooled samples were injected and analyzed using an Acquity UPLC Plus coupled through an electrospray ionization source to a Waters Synapt XS. Like serum samples, ions were separated using a Cogent Diamond Hydride HILIC chromatography column (2.2 µM, 120 Å, 150 mm x 2.1 mm, MicroSolv Leland, NC, United States) at a flow rate of 0.400 uL/min.

Pooled LC-MS/MS data were analyzed using Progenesis QI (Nonlinear Dynamics, Newcastle, UK, version 3.0). Data were imported, peaks were determined, and spectra were aligned. Next, acquired parent and daughter fragments were compared against theoretical fragmentation patterns using the Human Metabolome Database(21) for metabolite identification. We defined successful metabolite identifications as those with a Progenesis score greater than 60/100 and a fragmentation score > 20. The properties that contribute to these scores include mass error, isotope distribution similarity, and retention time error. Parts per million (ppm) error was calculated between LC-MS and LC-MS/MS data, and those with a ppm error greater than 20 were not considered.

## Results

### Changes in BMI and weight after 18-months of intensive weight loss interventions

Both changes in weight and BMI over 18 months were calculated for all participants selected for this analysis (n=30, n=10 per intervention). At baseline, 96.7% of participants selected for metabolomics had a BMI of 30 or greater. At 18 months, 62.5% of D+E participants, 40.0% of D participants, and 22.2% of E participants had a BMI less than 30. Considering weight loss, D+E (↓11.8%, p = 0.012) and D (↓9.4%, p = 0.0075) participants lost more weight than the E group (↓3.9%, p = 0.2973) (Supplementary Table 1)(3, 4). Weight loss and BMI measures for participant selected for this analysis were representation of the overall IDEA population (n=454)(4). Beyond the average weight loss per group, the amount of weight loss independent of group assignment was examined. Specifically, ranges of weight loss were categorized into 0-5%, 5-10%, 10%+ and weight gained and was determined to not influence metabolic results. However, sample size may limit conclusions and requires investigation among a larger subset of IDEA participants. Of the participants selected for this analysis, one E participant had a net weight loss of zero, and one participant from the D and D+E groups gained weight over 18 months. Weight loss data was not available for four participants in the randomly selected participants for this analysis (Supplementary Table 1).

### Weight loss interventions differentially influence the serum metabolome over 18 months

To examine variations in metabolism over the course of the 18-month period in response to three interventions – D, E, D+E – we calculated the changes between pre-trial and 18-month metabolite feature intensities, termed delta change (e.g., for each co-detected metabolite we subtracted the pre-trial intensity from the intensity at 18 months on a per-participant basis). The calculated delta change values were used for analysis throughout this study, for both intervention- and sex-associated differences. Across all intervention groups 2,142 unique metabolite features were co-detected. PCA and PLS-DA were used to analyze global metabolic differences in the calculated delta change values across the three intervention groups: diet (D), exercise (E), and combined diet and exercise (D+E).

PCA displays some overlap between groups with principal components (PC) 1 and 2 representing 32.3% of the total variability between groups (Figure 1A). PLS-DA shows greater separation and minimal overlap of groups with components 1 and 2 accounting for 5.2% and 12% of the variability, respectively (Figure 1B), suggesting that changes in metabolism reflect intervention type. A median metabolite intensity heatmap was used to perform additional analysis on the metabolite features with the 25 highest VIP scores (Figure 1C). VIP scores are calculated by summing the squared correlations between PLS-DA components and the original value and the metabolite feature. Of these 25 metabolites, 11 were more abundant in D participants, whereas 13 were more abundant in E participants and 1 was more abundant in D+E participants. Interestingly, the metabolite feature intensities in the D+E group overlap with those detected in the D and E group. The heatmap analysis of the top 25 VIP metabolite features demonstrated substantial metabolic differences between intervention types (Figure 1C).

**Figure 1.**
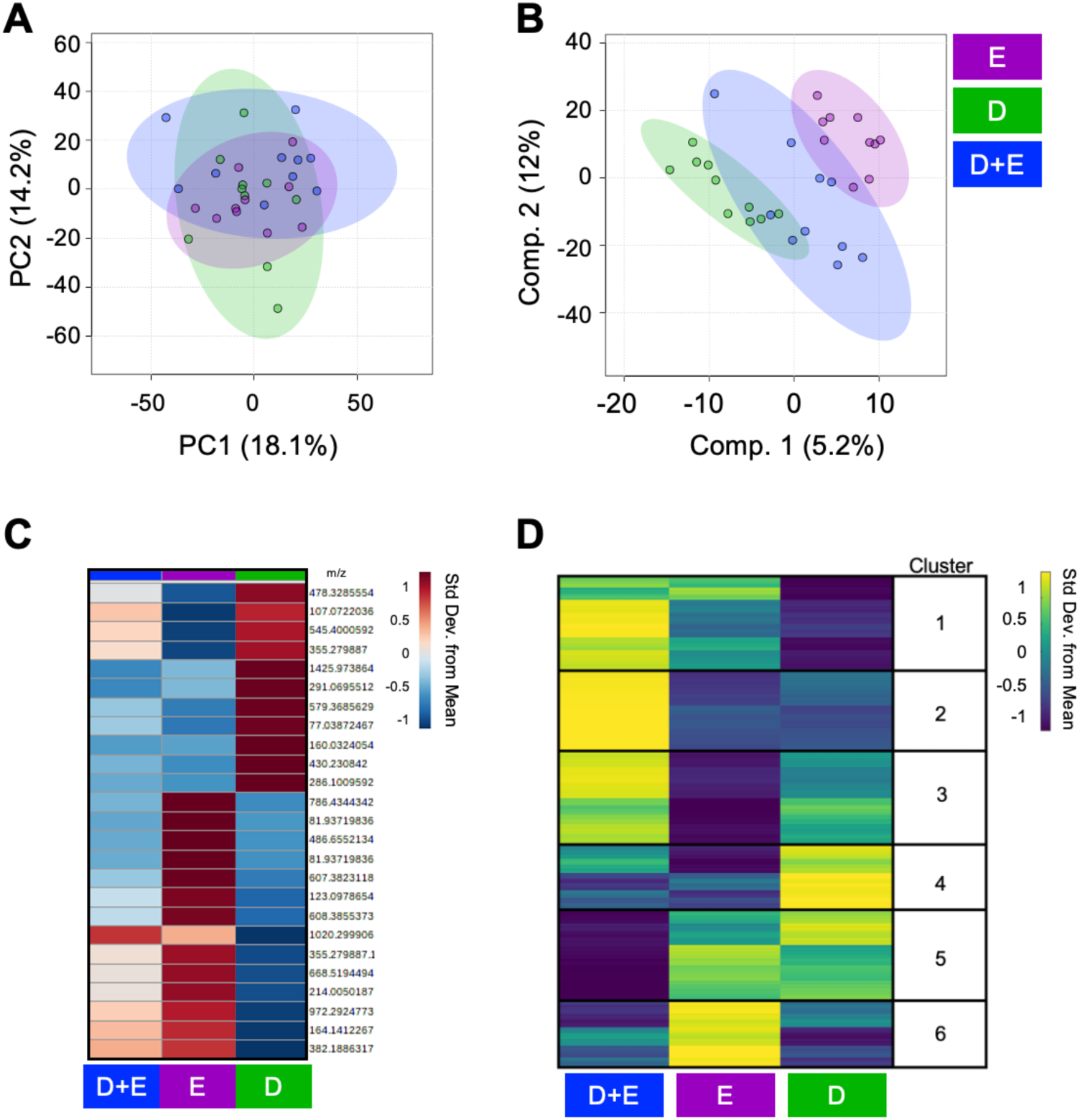
Metabolomic profiles differ between intervention groups after 18-months. (A) Principal component analysis (PCA), an unsupervised multivariate test, displays overlap of intervention groups in 2D, whereas (B) Partial least squares-discriminant analysis (PLS-DA), a supervised multivariate test, displays less overlap of interventions suggesting that the metabolome of participants assigned to different intervention groups for 18-months are distinct from each other. (C) A group median heatmap analysis of the top 25 PLS-DA Variable in Importance Projection (VIP) Scores, visualized using hierarchical clustering, highlights that exposure to 18-months of different interventions are associated with distinct metabolic regulation patterns. (D) Clusters of coregulated metabolite features (1–6) derived from participants serum across interventions underwent pathway enrichment analysis revealing distinct metabolic patterns that are associated with intervention type. Cooler colors (blue) and warmer colors (yellow) indicate lower and higher metabolite intensities relative to the mean. Colors in A-D correspond to: green – diet (D); purple – exercise (E); blue – diet and exercise (D+E).

To assess pathways affected by these interventions, we performed functional pathway enrichment analysis. First, we created a median metabolite intensity heatmap using delta values to distinguish clusters of similarly or differentially regulated metabolites on a global scale (Figure 1D). These clusters were analyzed using MetaboAnalyst’s Functional Analysis feature to underpin any significant pathways (false discovery rate adjusted p-values ≤ 0.05). Metabolite features highest in abundance among D+E participants and lowest in D participants mapped to xenobiotics metabolism, R-group synthesis, various fatty acid metabolisms, vitamin E metabolism, carnitine shuttle-related metabolism, and cytochrome P450 metabolism. Comparatively, metabolite features highest in D+E participants and lowest in both D and E groups mapped to di-unsaturated fatty acid beta-oxidation, linoleate metabolism, dynorphin metabolism, glycerophospholipid biosynthesis and metabolism, and squalene and cholesterol biosynthesis. Features highest in D+E participants and lowest in the E participants mapped to polyunsaturated fatty acid biosynthesis, glycerophospholipid metabolism, biopterin metabolism, alkaloid biosynthesis, and fatty acid activation and oxidation.

Metabolite features highest in abundance among D participants and lowest in both the D+E and E participants mapped to metabolism of vitamin metabolism (B6, C), carbohydrate and sugar metabolism (hexose, starch, sucrose, fructose, mannose), lipid-related pathways (glycerophospholipid, glycosphingolipid, omega-6), N-glycan biosynthesis and degradation, as well as pathways related to energy metabolism (pentose phosphate, glycolysis, gluconeogenesis). Metabolites features highest in abundance among E participants and lowest in D+E and D groups mapped to amino group metabolism (glutamate, histidine, methionine, cysteine, tryptophan, alanine, and aspartate), nitrogen metabolism, purine metabolism, vitamin B3 metabolism, and metabolisms involved with the formation of putative anti-inflammatory metabolites from eicosapentaenoic acid. Finally, metabolites highest in both E and D groups and lowest in D+E participants mapped to various amino group metabolisms (glycine, serine, threonine, arginine, proline, alanine, aspartate, lysine, and asparagine), N-glycan degradation, urea cycle, butanoate metabolism, chondroitin sulfate degradation, keratan sulfate degradation, heparan sulfate degradation, and cytochrome P450 metabolism (Table 2).

**Table 2.**
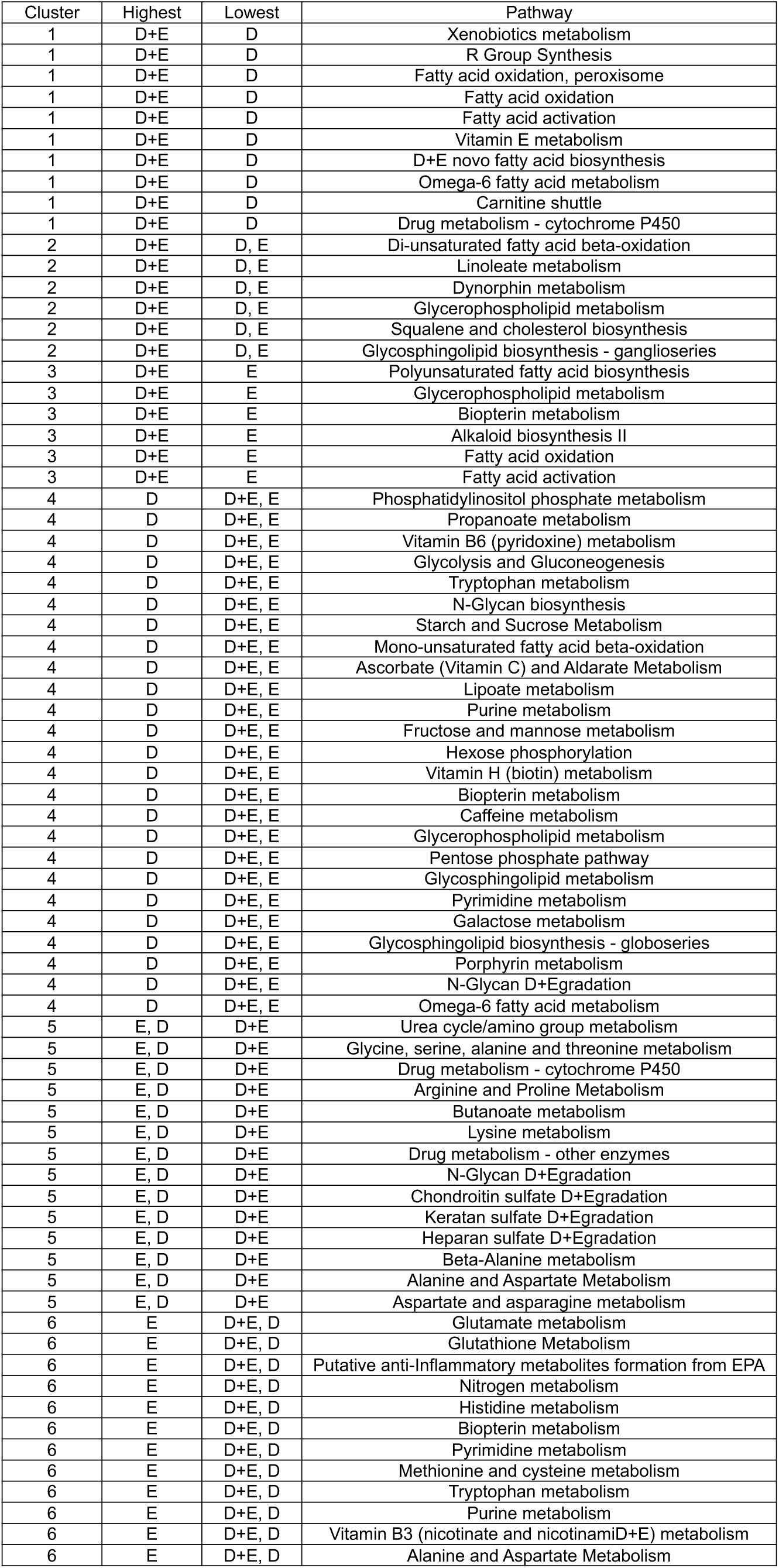
Metabolic pathways associated with diet, exercise, and diet plus exercise interventions identified by median metabolite intensity heatmap analysis. All reported pathways have a FDR-corrected significance level < 0.05. D = diet. E = exercise. DE = diet and exercise. Clusters defined in Figure 1D.

Additionally, pairwise comparisons using delta change values between the three intervention groups were performed to further investigate how weight loss interventions differentially influence OA metabolism (Supplementary Figure 2, Supplementary Table 2). Specifically, PCA and PLS-DA were used to visualize the metabolomes of participants and is detailed in supplementary results. Populations of metabolite features distinguished by fold change when comparing intervention groups underwent pathway analyses. Pathways associated with both D+E and E groups included N-glycan biosynthesis and ubiquinone and other terpenoid-quinone biosynthesis. All other overlapping pathways were associated with both D and E participants which included aminoacyl-tRNA biosynthesis, cytochrome P450 metabolism, purine metabolism, and glutathione metabolism. No significant pathways were associated exclusively with either the D or E participants (Supplementary Table 3). Additionally, populations of metabolite features distinguished by fold change when comparing intervention groups were matched against metabolite identifications made using LC-MS/MS data. Many of these were detected across comparisons and displayed diverse regulation patterns (Supplementary Table 4). Altogether, the results show that different weight loss interventions influence OA metabolism in distinct ways.

### Serum metabolome is affected by sex after 18-months of intervention

To examine potential sexual dimorphism within the serum metabolome, PCA, PLS-DA, fold change, and volcano plot analyses were used to compare male and female participants, both in general and within intervention groups. PCA comparison of all male and female participants finds overlap with principal components 1 and 2 accounting for more than 30% of the variability in the dataset. PLS-DA reveals distinct endotypes based on sex, with components 1 and 2 accounting for 19.6% of the variability (Figure 2A-B). Fold change revealed that 283 metabolites had a greater change (FC>2) in abundance during the 18-month period in female participants than in male participants, mapping to glycolysis/gluconeogenesis, ubiquinone and other terpenoid-quinone biosynthesis, various amino acid metabolisms, and metabolism of starch and sucrose. Comparatively, 345 features had a greater change in abundance during the 18-month period in male participants than female participants, and these metabolites mapped to purine metabolism (Figure 2C, Supplementary Table 5).

**Figure 2.**
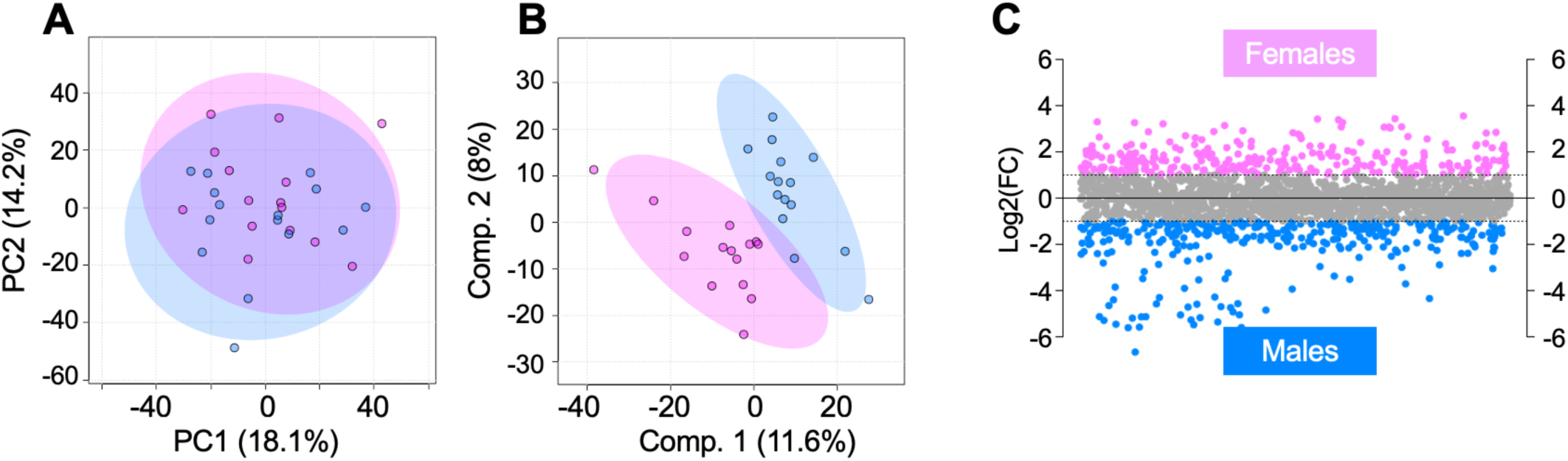
Serum metabolome of all IDEA participants, excluding the factor of intervention, is influenced by participant sex. (A) While PCA displays overlapping of male and female IDEA participants, (B) PLS-DA improves clustering of participants within their respective sex group suggesting the serum metabolome is sexually dimorphic. (C) Fold change analysis was conducted to pinpoint populations of metabolite features that are differentially regulated between males and females. This revealed 283 metabolite features that had a FC > 2 and were higher in abundance in female participants, whereas 345 had a FC < -2 and were higher in abundance in male participants. The colors in A-C correspond to: pink – females; blue – males.

Likewise, a pairwise comparison was conducted between male and female D+E participants. PCA finds some overlap of male and female participants, with PCs 1 and 2 accounting for 47.3% of the total variability (Figure 3A). PLS-DA, however, finds complete separation of participants based on sex with components 1 and 2 representing 26% of the variability (Figure 3B). Among female participants, 586 metabolites had at least a 2-fold greater change in abundance during the 18-month period than they did in male participants. These metabolites mapped to tryptophan metabolism, glycosaminoglycan degradation, biosynthesis of unsaturated fatty acids, and caffeine metabolism. 532 features had at least a 2-fold increase in male participants. These mapped to aminoacyl-tRNA biosynthesis, lysine degradation, various amino acid metabolisms, and glutathione metabolism (Figure 3C, Supplementary Table 5).

**Figure 3.**
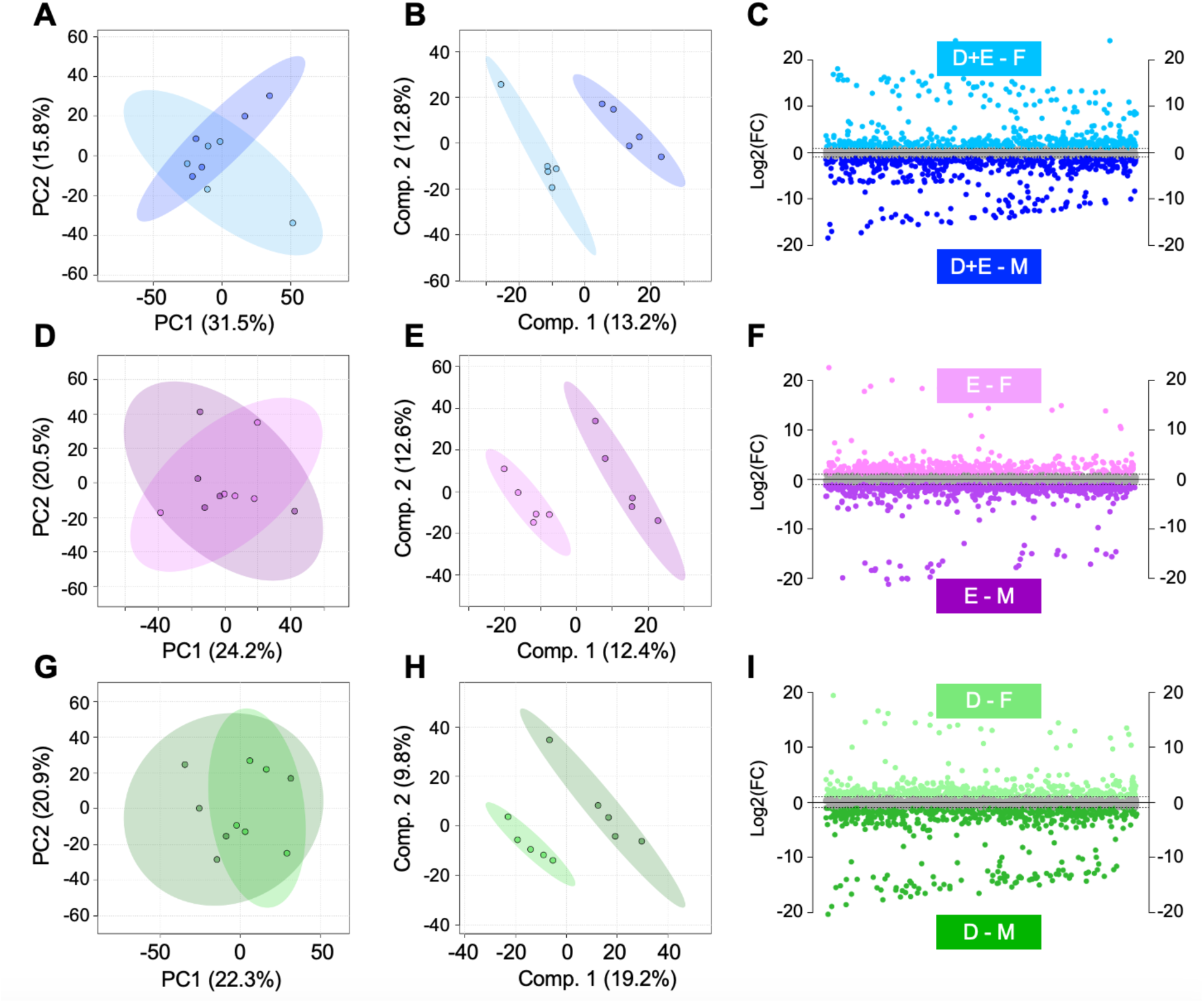
Global analysis of intervention groups after 18-months reveals that the serum metabolome is influenced by sex. (A) PCA displayed some overlap of D+E males and females whereas (B) PLS-DA perfectly separated D+E males and females suggesting that the serum metabolome is potentially influenced by participant sex. (C) Fold change analysis identified 568 metabolite features that had a FC > 2 and were higher in abundance in D+E females. Conversely, 532 had a FC < -2 and were higher in abundance in D+E males. Similarly (D) PCA, (E) PLS-DA, and (F) fold change was applied and revealed a similar sexual dimorphic pattern when comparing male and female E participants. This same suite of analyses was applied to examine sex-associated metabolic patterns between male and female D participants (G-I). Populations of metabolite features distinguished by fold change analyses were then subjected to functional pathway enrichment analyses to pinpoint biological pathways. Considering differences in metabolic regulation associated with sex across D+E (A-C), E (D-F) and E (G-I) participants, it is evident that that serum metabolome of IDEA participants is influenced by sex. The colors in A-I correspond to: light and dark blue – female and male diet and exercise (D+E) participants; light and dark purple – female and male exercise (E) participants; light and dark green– female and male diet participants.

Among E participants, PCA shows some overlap between male and female participants, with PCs 1 and 2 accounting for 44.7% of the total variability (Figure 3D). PLS-DA shows complete separation of groups with components representing 25% of the variability (Figure 3E). Among female participants, 433 metabolites had a 2-fold or greater change in abundance than they did in male participants, mapping to caffeine metabolism. 617 metabolite features had a greater change in male participants compared to female participants, mapping valine, leucine, and isoleucine biosynthesis (Figure 2F, Supplementary Table 5).

Finally, in D participants, PCA shows overlap between male and female participants, with the first two PCs accounting for 43.2% of variability in the dataset (Figure 2G). PLS-DA shows complete separation of groups with components representing 29% of the variability (Figure 2H). Additionally, 524 metabolites had a 2-fold or greater change in abundance in female participants than they did in male participants. These mapped to aminoacyl-tRNA biosynthesis, and valine, leucine, and isoleucine biosynthesis. 632 features had a greater change in male participants, mapping to purine metabolism (Figure 2I, Supplementary Table 6).

Populations distinguished through fold change analyses assessing sexual dimorphism across intervention groups (Figure 3) were matched against LC-MS/MS identified metabolites. Notably, many of these identified metabolites were statistically significant across pairwise comparisons between interventions and were sex-associated. Of these, docasanedioic acid showcased consistently higher abundance patterns in all female participants within each intervention groups. Excluding D male and female participants, stearoylcarnitine was higher in abundance in female participants when comparing metabolic patterns associated with sex within the E and D+E groups. Strikingly, N-acetyldemethylphosphoinothricin was detected when investigating both intervention- and sex-associated differences. Specifically, it was higher in male participants when examining sex differences within intervention groups. However, when considering interventions only, it was the highest among D participants compared to E and D+E participants (Supplementary Table 4). In summation, the data vividly illustrates the striking divergence in serum metabolism between male and female participants subjected to different weight loss interventions.

## Discussion

This metabolomics study, in conjunction with the IDEA trial, demonstrates that intensive weight loss intervention, including diet and combined diet and exercise, influences the metabolome of OA patients differently from exercise. When comparing participants after 18-months of different weight loss interventions, changes in energy, vitamin, carbohydrate, amino acid, and lipid metabolism were detected. Moreover, these data demonstrate clear metabolic differentiation between IDEA participants based on participant sex. Overall, intensive weight loss and exercise influence the serum metabolome of OA individuals, and sex may independently influence the nature of metabolic changes in OA. The integration of the present metabolomic profiles with the previously reported IDEA study outcomes(4, 17) enhances the understanding of the effects of dietary weight loss and exercise in individuals with OA and provides insights into intervention effectiveness and potential biomarker identification.

Previously, NMR metabolomics was used to assess urine from IDEA participants finding detected metabolites corresponding to energy, amino acid, and lipid metabolisms. The present study detected similar metabolic themes in serum extracts across intervention groups. D+E participants exhibited elevated lipid-related pathways such as the carnitine shuttle compared to both D and E participants (Table 2). Notably, the carnitine shuttle’s primary function is facilitating transport of fatty acids into the mitochondria for ATP generation via beta-oxidation. A prior study finds that short-term consumption of a high fat diet increases levels of circulating fatty acids and carnitines(22). Specific to OA, higher concentrations of carnitines are observed in synovium from end-stage knee OA patients(23). Additionally, L-carnitine supplementation is a potential treatment to mitigate OA-related inflammation and oxidative stress(24, 25).

D+E participants also displayed changes in other noteworthy lipid-related pathways such as polyunsaturated fatty acid (PUFA) biosynthesis, omega-6 fatty acid metabolism, linoleate metabolism, and glycerophospholipid metabolism after 18-months of intervention. Omega-3 and -6 fatty acids, as well as linoleic acid, a polyunsaturated omega-6, have all been implicated in the pathogenesis of OA(26, 27). Glycosphingolipids, a key membrane component, have been previously linked to exercise, obesity, metabolic syndrome, and insulin resistance(28–30). Dysregulation of membrane-related pathways may stem from exercise intervention (E and D+E participants), requiring further investigation. These findings suggest that the combination of diet and exercise act on pathways like the carnitine shuttle, PUFA, and glycosphingolipid metabolism to impact fat utilization for energy, oxidative, and inflammatory functions.

D participants had elevated glycolysis, pentose phosphate pathway, purine and pyrimidine metabolism, and various sugar and vitamin metabolisms compared to E and D+E participants (Table 2). Glycolysis and the pentose phosphate pathway play central roles in central energy metabolism as well as production of nucleotides and amino acid precursors, oxidative stress reduction, and ATP generation. Disruptions in these pathways can lead to insufficient ATP, NADH accumulation, inflammation, generation of reactive oxygen species, and altered rates of gluconeogenesis.

Carbohydrate pathways like starch and sucrose metabolism are linked to central energy metabolism, possibly serving as biofuels for ATP production. Notable related pathways are hexose phosphorylation and vitamin C metabolism which were both detected among D participants. Vitamin C is a hexose derivative from glucose that has antioxidant capabilities and serves as a defense against reactive oxygen species, is critical for collagen synthesis by enhancing pro-collagen hydroxylation and has been hypothesized to have a protective role in OA(25, 31, 32). Moreover, a study that investigated the association between vitamin C and knee OA found that low intake of vitamin C is a potential risk factor for KOA(32). However, neither vitamin intake nor serum concentration was measured in this study. Additionally, likely variability in participant meal and snack choices warrants further investigation to understand the relationship between vitamin and carbohydrate metabolism, diet intervention, and OA.

Exercise is a common treatment for improving symptoms of obesity and related conditions such as cardiovascular disease, type II diabetes, and OA(33, 34). Additionally, the anti-inflammatory effects of exercise have been well documented(35–37). This could explain the detection of increased putative anti-inflammatory and amino acid pathways including glutamate, histidine, alanine, aspartate, methionine, cysteine, and glutathione metabolism across E participants compared to D+E and D participants. Thus, the finding that these pathways were highest in E participants and detected in lower abundances in both D and D+E participants suggests that these metabolic changes are as a result of the diet. Notably, cysteine and glutamate are essential to produce glutathione, a key antioxidant that helps scavenge reactive oxygen species produced during normal cell metabolism and has been hypothesized to play a key role in the inflammatory response in OA(38). Oxidative status, as reflected by the ratio of cysteine, glycine, and glutamate, tends to decrease with age, thereby increasing stress and cell death in chondrocytes(39). Therefore, detection of glutathione, cysteine, and glutamate metabolism may reflect oxidative stress resistance in the presence of exercise. Moreover, amino acids like alanine, aspartate, and glutamate have been previously associated with OA and have been detected in higher concentrations among OA patients(40). Histidine concentration distinguishes between the synovial fluid of OA and RA patients(41) and decreases as OA progresses(42). Taken together, these findings suggest that metabolism of these amino acids may undergo significant changes in response to exercise in OA patients, suggesting amino acids could potentially be monitored overtime to oversee OA development. Moreover, additional research is needed to better understand the relationship between amino acid metabolism, weight loss interventions, and OA.

While it is recognized that knee OA is more common and often more severe in female patients(5, 6), metabolic differences associated with knee OA that differ by sex remain uncertain. An increasing number of studies find sexual dimorphism in the context of OA and contribute mounting evidence indicating that the etiology and progression of OA may exhibit sex-specific changes as shown in these data. Divergent pathways between male and female IDEA participants mapped to glycolysis/gluconeogenesis, amino acid metabolism, and lipid-related metabolisms (Supplementary Table 5).

Sex-dependent differences in bone size and angle and pelvis and joint size are well documented(43, 44). At the hormonal level, females have greater circulating levels of estrogen with levels fluctuating throughout the menstrual cycle and post-menopause. Thus, it is not surprising that these varying hormone levels appear to influence the serum metabolome. Both E and D+E female participants had elevated caffeine metabolism compared to male participants. For pre- and post-menopausal women, hormonal cycles may affect amino acid metabolism where protein oxidation is impacted by estrogen levels(45). Understanding these sex-related metabolic differences could refine current OA prevention and treatment strategies.

This study is not without limitations. First, this study lacks serum from both healthy controls and individuals with OA who did not undergo an intervention. Second, serum-derived metabolomic data may reflect joint health and provide insight into the effects of intervention and sex. However, the relationship between serum and synovial fluid is complex with many metabolites correlated between these compartments but many more metabolites without correlations(46, 47). As such, the relationship between joint health and the serum metabolomic requires further investigation. A third limitation is the relatively small samples size of 10 individuals per intervention group which was necessitated by the time and cost of performing an extensive metabolomics analysis.

The outcomes of this comprehensive metabolomics investigation, in conjunction with the overarching findings of the IDEA study, emphasize that intensive weight loss has symptomatic and potential metabolic benefits among individuals with knee OA. Metabolic perturbations associated with these interventions, as well as patient sex, likely play a role in OA development and should be considered for downstream treatment strategies. By further applying metabolomics in parallel with these findings a greater understanding of the joint-level dysregulation during OA progression may shed light on the impacts of obesity, weight reduction, patient treatments, and interventions. Expansion with larger sample sizes will delve into the associations between other measurements obtained throughout the duration of the IDEA study and metabolomic results among all study participants. These insights hold promise for enhancing the current approach to managing and improving the quality of life of those affected by OA.

## Acknowledgements

We thank the Montana State University Mass Spectrometry Facility including Dr. Donald Smith and Jesse Thomas for aiding in LC-MS analysis and interpretation. Funding for the Montana State University Mass Spectrometry Facility was made possible through the M.J. Murdock Charitable Trust, the National Institute of General Medical Sciences of the National Institutes of Health (P20GM103474 and S10OD28650). Additionally, authors thank Brady Hislop for his assistance in developing analysis pipelines and analyzing data. This study was supported by grants from the National Institutes of Health (NIAMS R01AR073964 and R01AR081489) and the National Science Foundation (CMMI 1554708).

## Author Contributions

HDW and RKJ designed metabolomics experiments; RFL and SPM designed trial parameters; HDW performed metabolite extractions; HWD ran LC-MS samples; HDW and AHW analyzed data; HWD, AHW, BB, STP, RFL, and RKJ interpreted results; HDW, AHW, and RKJ drafted manuscript. All authors have read and revised the manuscript.

---

**Supplementary Figure 1.**
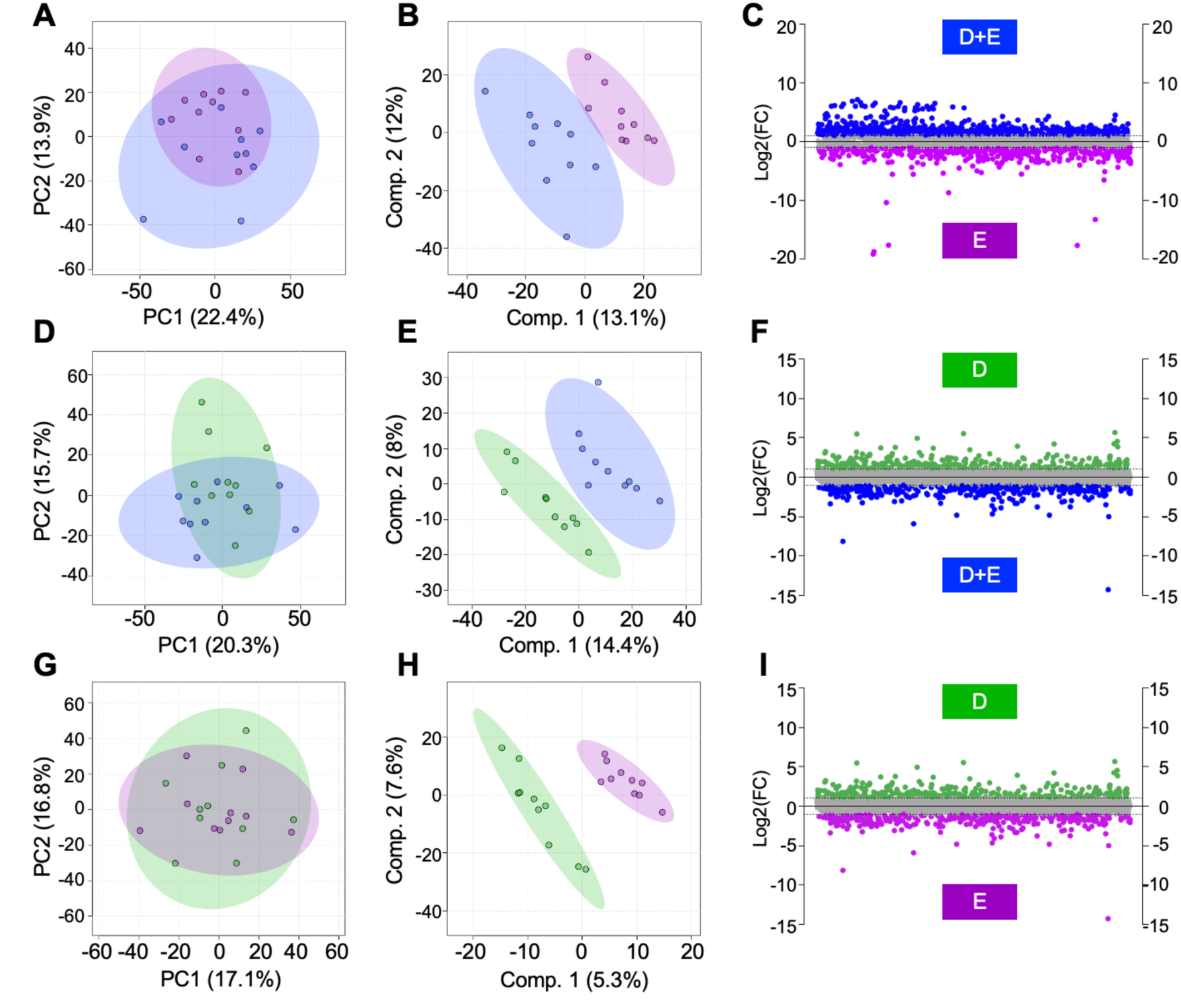
Pairwise comparison unveils intervention-associated metabolic patterns. (A) Principal component analysis and (B) partial least squares-discriminant analysis were applied to visualize the metabolome of D+E and E participants after 18-months of intervention. (C) Fold change analysis identified 760 metabolite features that had a FC > 2 and were higher in abundance in D+E participants. Conversely, 429 had a FC < -2 and were higher in abundance in E participants. Similarly, (D) PCA displayed some overlap, whereas (E) PLS-DA displayed clear separation of D+E and D participants. (F) Fold change analysis was applied to examine metabolic differences associated with D+E and D participants. This same suite of analyses was applied to investigate metabolic differences between E and D participants (G-I). Populations of metabolite features distinguished by fold change analyses were then subjected to functional pathway enrichment analyses to pinpoint biological pathways. Collectively, it is evident that the serum metabolome is influenced by intervention type, and differences in metabolic patterns reflect intervention status. The colors in A-I correspond to: green – diet (D); purple – exercise (E); blue – diet and exercise (D+E).

